# Neurotropic strains of *Listeria monocytogenes* preferentially invade enteric glial cells

**DOI:** 10.64898/2026.08.03.742483

**Authors:** Ryan W. Donkin, Caroline F. Benda, Katelynn E. Krick, Breanna Amelunke, Jooyoung Cho, Ellie L. Sams, Taylor M. Albrecht, Angélica M. Peña Rosado, Taylor E. Senay, Anna C. Puderbaugh, Joshua S. Nowacki, Sarah E.F. D’Orazio

## Abstract

Certain strains of the facultative intracellular bacterial pathogen *Listeria monocytogenes* are thought to invade cranial nerves in the gut and disseminate directly to the brainstem to cause rhombencephalitis in both humans and ruminants. Bacteria with actin tails were previously observed within neurons of naturally infected sheep, but the mechanism for how these neurotropic strains access the nervous system has not been well characterized. Using a foodborne mouse model of listeriosis, we show here that bypassing the gut phase of infection prevents colonization of the brain, confirming that invasion of the nervous system happens in the intestinal tract. *L. monocytogenes* did not efficiently invade neuronal cell lines, although they could replicate exponentially in the cytosol and form actin tails. Instead, the neurotropic strains displayed a preferential ability to invade enteric glial cells, a specialized subset of glia that support neurons and are critical for intestinal homeostasis. Using an in vitro co-culture system, we demonstrated that neurotropic *L. monocytogenes* could readily invade enteric glial cells and use ActA-mediated actin-based motility to spread to adjacent neurons. These results suggest that invasion of enteric glial cells is a novel virulence strategy that can promote brainstem infection following foodborne transmission of *L. monocytogenes*.

**IMPORTANCE:** This study provides further evidence for dissemination of neurotropic strains of *L. monocytogenes* from the gut directly to the brain via axonal migration using foodborne mouse model of listeriosis. It is the first report showing that enteric glial cells, a specialized subset of cells in the gut that support intestinal neurons, are susceptible to pathogenic bacterial infection.

## INTRODUCTION

*Listeria monocytogenes* is a Gram-positive facultative intracellular bacterium that causes foodborne illness in humans. Research investigating the pathogenesis of listeriosis has focused largely on reference strains EGDe and 10403s, but in recent years, many novel virulence genes have been discovered by doing comparative genomic analyses amongst *L. monocytogenes* isolates (1). Four phylogenetic lineages of *L. monocytogenes* have been described, with the majority of sequenced isolates belonging to either lineage I or II (2). Much less is known about strains in lineages III and IV, which seem to be less common(3). In this study, we explored the neurotropism of two lineage III strains: UKVDL9, which was isolated from a sheep brain and SD4000, which was isolated from a patient with a brainstem infection (rhombencephalitis) (4).

It has been proposed that *L. monocytogenes* can use one of three mechanisms to enter the brain. The bacteria can either directly invade endothelial cells in the blood-brain barrier (5) or be transported while associated with circulating monocytes across the barrier (6) and then disseminate diffusely to cause meningoencephalitis. Both of these mechanisms require high titer bacteremia, and therefore, are associated with either immune compromise or a very large inoculum (7, 8). The third strategy involves invasion of cranial nerves in the gastrointestinal tract and ActA-mediated motility along the axon to the brainstem, resulting in a focal brainstem infection known as rhombencephalitis. These infections can occur in otherwise healthy humans (9), suggesting that the causative bacterial strains have a tropism for invading the nervous system. Rhombencephalitis is the most common form of listeriosis in ruminants, and it has been suggested that cows and sheep may serve as a reservoir for these neurotropic strains (10). We previously showed that rhombencephalitis isolates UKVDL9 and SD4000 could disseminate to the brain following foodborne transmission in mice and cause both acute and lingering neurological deficits (11). In contrast, neither reference strain EGDe nor other lineage III strains we tested disseminated to the brain in the murine natural feeding model.

*L. monocytogenes* can be readily internalized by macrophages or can induce its own uptake when surface exposed internalin proteins engage host cell receptors on epithelial cells or endothelial cells (12). Most of the bacteria are thought to escape from the phagocytic or endocytic vacuole due to the actions of the pore-forming toxin listeriolysin O and two phospholipases (13). The surface protein ActA undergoes a polar redistribution once the bacteria begin replicating in the cytosol and this enables the bacteria to co-opt the host cell actin polymerization machinery to promote actin-based motility, a type of intracellular movement (14, 15). Direct cell-to-cell spread can occur when actin-based motility drives the formation of protrusions which can be resolved into double membrane vacuoles in the neighboring cell if they occur in the presence of both the bacterial and host cell proteins needed to relieve membrane tension and promote internalization (16). This intracellular life cycle is extremely well characterized in macrophages. Escape to the cytosol typically happens within 30 minutes, actin tails begin to appear within a few hours once the bacteria start replicating, and cell-to-cell spread is observable as foci of infection in a monolayer of cells within several hours. Much less is known about the parameters of this intracellular life cycle in other cell types.

In this study, we provide further evidence that the dissemination of neurotropic strains of *L. monocytogenes* to the brain following foodborne transmission requires intestinal colonization rather than originating from growth in systemic tissues such as the spleen and liver. We surprisingly showed that *L. monocytogenes* were not able to efficiently invade neurons, but the few bacteria that are internalized can replicate exponentially in the cytosol of neurons. Instead, we found that the key advantage of neurotropic strains compared to reference strain EGDe was an enhanced ability to be internalized by enteric glial cells, a unique and important but understudied cell type in the gut that support neurons and promote intestinal homeostasis. Importantly, we showed that *L. monocytogenes* were able to use ActA to spread from an enteric glial cell into the cytosol of a neuron, providing a mechanism for the bacteria to dissemination from the gut directly to the brainstem without reaching high titer in the blood.

## RESULTS

### Neurotropic strains UKVDL9 and SD4000 do not overexpress InlB

Direct invasion of the blood-brain barrier by *L. monocytogenes* is thought to involve interactions between the bacterial cell surface protein internalin F and host cell vimentin (5); however, we previously showed that neurotropic strains UKVDL9 and SD4000 both lack an internalin F ortholog similar to other lineage III strains (3, 11). Maudet et al. recently reported an alternate strategy to promote invasion of the blood brain barrier which requires InlB-dependent signaling through c-Met to inhibit Fas-mediated killing of infected monocytes by T cells, allowing the monocytes to carry *Listeria* across the barrier in a “Trojan horse” style (17). In that study, certain hypervirulent strains rapidly invaded the brain within 2 days of intravenous infection; these strains all expressed four-fold increased levels of *inlB* mRNA relative to *L. monocytogenes* reference strains EGDe and 10403s.

To determine if our neurotropic *L. monocytogenes* strains expressed the increased level of *inlB* needed to promote monocyte-mediated invasion of the blood-brain barrier, we assessed *inlB* mRNA levels in late logarithmic phase cultures. In three independent experiments, there was no significant difference in *inlB* expression for strain UKVDL9 compared to EGDe and strain SD4000 expressed significantly less *inlB* mRNA than the reference strain (Fig 1A). We were unable to test the specific clonal complex 1 (CC1), CC4, and CC6 strains used in the Maudet et al. study because of institutional disagreements arising from the proposed Materials Transfer Agreement. However, we tested two other *L. monocytogenes* strains known to be hypervirulent in the intravenous model of listeriosis (PF49 and P14) (18)and found that they also did not have increased *inlB* expression (Fig. 1A). These data are consistent with our hypothesis that strains UKVDL9 and SD4000 disseminate to the brain without needing to persist in the bloodstream (11) and without using InlB-dependent mechanisms to cross the blood-brain barrier.

**FIG 1.**
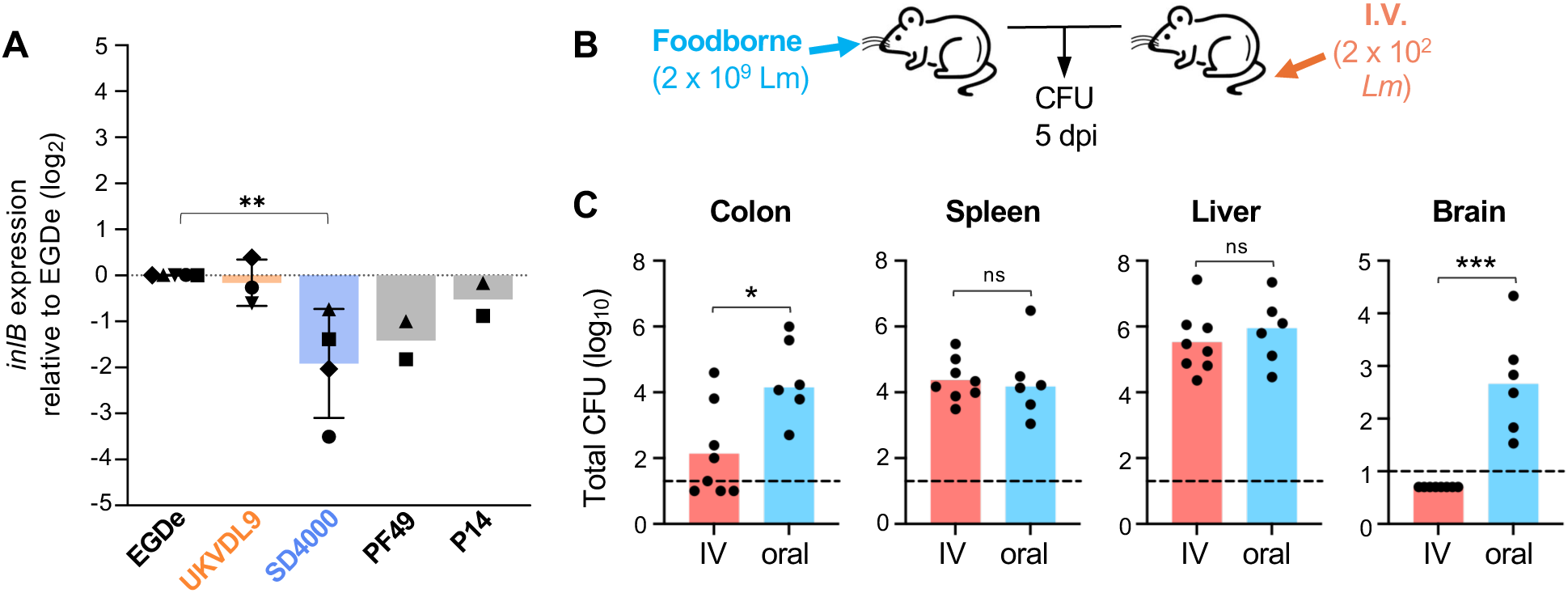
Neurotropic *L. monocytogenes* disseminate to the brain only when the transmission route involves the gastrointestinal tract. (A) The indicated *L. monocytogenes* strains were grown to late log phase and *inlB* mRNA expression was measured by qRT-PCR. Each symbol represents the mean 2^-ΔΔCt^ value of *n*=3 technical replicates normalized to the mean EGDe value for that experiment (RNA was harvested in five separate experiments indicated by the shape of the symbol). Bars indicate the mean 2^-ΔΔCt^ values (± SD) across independent experiments. Statistical significance was assessed by one-way ANOVA with Dunnett’s multiple comparisons test. **(B)** Female BALB/cByJ mice were either fed (oral) 2 x 10^9^ CFU of *L. monocytogenes* (Lm) SD4000 or injected intravenously (IV) with 2 x 10^2^ CFU of the same strain. **(C)** Tissues were harvested 5 days post-infection for CFU determination; dashed lines indicate the limits of detection. Symbols indicate CFU values from individual mice; pooled data from two independent experiments are shown. Bars indicate median values which were analyzed by Mann-Whitney test.

### Dissemination to the brain requires colonization of the intestines

To more directly test whether invasion of the nervous system happened during the intestinal phase of infection or after *L. monocytogenes* had spread systemically, we assessed colonization of the brain following foodborne or intravenous (i.v.) infection. Foodborne infection requires a much higher inoculum than i.v. administration because only a fraction of the ingested bacteria survive passage through the stomach while nearly all of an i.v. inoculum can be detected in the spleen and liver within 15 minutes (19–21). Both groups of mice received a sublethal dose appropriate for the transmission route (Fig. 1B), and at five days post-infection equivalent bacterial burdens were observed in the spleen and liver (Fig. 1C). Oral infection resulted in higher bacterial burdens in the colon than did i.v. infection; this was expected since colonization of the intestines following i.v. inoculation is thought to occur only late in the infection after the gallbladder becomes colonized and infected bile is released into the gut (19, 22). Most notably, foodborne infection resulted in substantia brain colonization, whereas none of the mice infected intravenously had *L. monocytogenes* in the brain (Fig. 1C). This suggests that bypassing the gut completely eliminated the opportunity for the neurotropic factor(s) encoded by strain SD4000 to promote dissemination to the brain.

### Co-localization of *Listeria* with neurons in the gut

The third mechanism for *L. monocytogenes* to disseminate to the brain does not involve the blood-brain barrier and instead happens when cytosolic bacteria use actin-based motility to spread within neurons directly to the brainstem resulting in rhombencephalitis. To look for association of *L. monocytogenes* with neurons in the gut, mice were fed neurotropic strains SD4000 or UKVDL9 or the reference strain EGDe and small intestines were harvested three days post-infection and stained with β-3-tubulin to identify neurons. Each intestine was rolled tightly and a cross section representing both the proximal and distal regions was examined microscopically (Fig. 2A). The number of *L. monocytogenes* visualized in the small intestines of each mouse varied from tens to hundreds of bacteria, but at least 920 distinct bacteria were observed in the pooled tissues infected with each strain (Fig. 2B). The percentage of *L. monocytogenes* found in close proximity to β-3-tubulin+ cells in four distinct anatomic regions of the gut tissue was determined (Fig. 2C, 2D).

**FIG 2.**
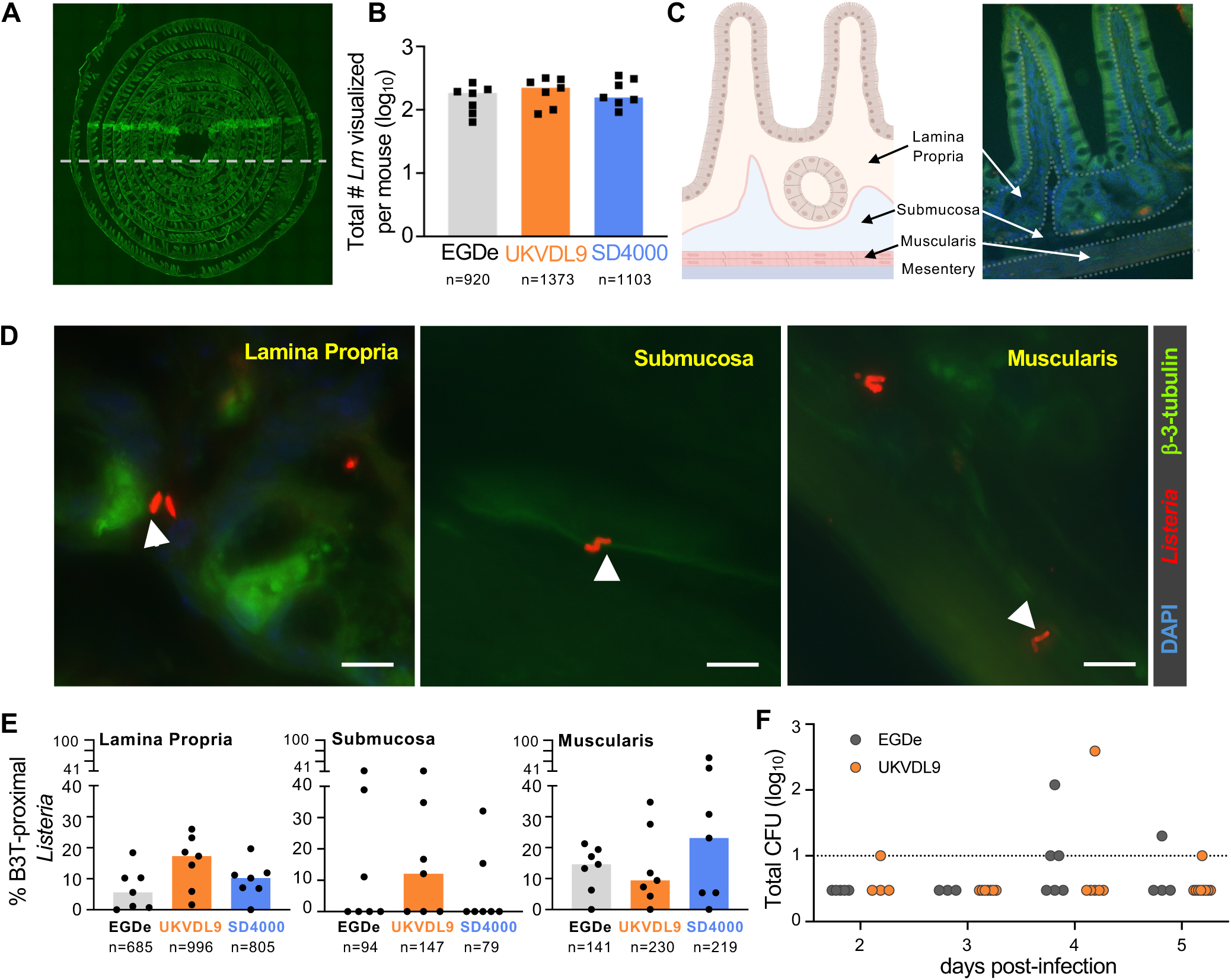
*L. monocytogenes (Lm)* were found in proximity to neurons in the small intestine. BALB/cByJ mice were fed ∼2 x 10^9^ CFU of *Lm* EGDe, UKVDL9 or SD4000 and small intestines were harvested 3 days post-infection. **(A)** Rolled tissue preps were fixed and stained and a cross section of each (dotted line) was visualized to locate bacteria ranging from the proximal to distal intestines. **(B)** Pooled data from two separate experiments (n=7 mice per group) were analyzed; symbols indicate the number of *Lm* observed in the intestines of individual mice. The total number of *Lm* analyzed in each sample group is indicated below the graph. **(C)** Observed bacteria were sorted into four anatomic regions indicated by the graphic on the left and the 40X image of one of the murine intestines on the right. **(D)** Representative images from each region show bacteria in proximity to beta-3-tubulin (B3T)-positive cells (white arrowheads). Scale bars, 10 µm. **(E)** The percentage of B3T-associated bacteria in each anatomic region is shown with symbols representing individual mice and bars indicating median values. The total number of *Lm* observed in each sample group is indicated below the graph. Data were analyzed by one-way ANOVA with Dunnett’s multiple comparisons test and no significant differences were observed. **(F)** Mice were fed 2 x 10^9^ CFU of *Lm* EGDe or UKVDL9 and the vagus nerve was harvested at the indicated timepoints for CFU determination. Pooled data from three separate experiments (n=4-9) are shown.

Most of the bacteria observed in each mouse were located in the lamina propria of the small intestine (Fig. 2E). Although there was a trend towards a higher percentage of the neurotropic *L. monocytogenes* in proximity to β-3-tubulin+ cells, the difference was not statistically significant. *L. monocytogenes* were also observed in proximity to neurons in both the submucosa and the muscularis, where the myenteric plexus of the enteric nervous system is found, but there were no differences between the three strains tested (Fig. 2E). Very few bacteria were observed in the underlying mesentery and none of them were in proximity to neurons (data not shown). These data suggested that foodborne transmission of *L. monocytogenes* results in multiple opportunities for the bacteria to interact with neurons in the gut, with highest probability of an interaction occurring in the lamina propria. However, this analysis supported a rejection of the hypothesis that strains UKVDL9 and SD4000 are neurotropic due to an enhanced ability to access neurons in the gut relative to reference strains like EGDe.

We next attempted to obtain more direct evidence of *L. monocytogenes* ascending to the brain via axonal migration by dissecting the vagus nerve and plating for total CFU. The vagus is the most prominent cranial nerve in the gut and it innervates the majority of the small intestine. We were able to recover 5 to 10 mm segments from the lower part of nerve near the esophageal sphincter, and used a Dounce homogenizer to process the tissue such that our limit of detection was only 10 CFU. We knew that *L. monocytogenes* did not colonize the brainstem until five days post-infection (11), but it was unclear whether the delay was due to initial invasion of the nervous system or the time needed to disseminate within the axon. Therefore, we harvested vagus tissue from groups of mice at 2, 3, 4, and 5 days post-infection. As shown in Fig. 2F, no *L. monocytogenes* were recovered from the majority of the mice we examined. Both EGDe and UKVDL9 were recovered, but the small number of animals with live CFU in the vagus precluded any statistical analysis. Thus, these data suggested that oral transmission of *L. monocytogenes* could result in infection of the vagus nerve, but did not indicate a particular timeframe when this was likely to occur.

### UKVDL9 and SD4000 inefficiently invade neurons

We next postulated that strains UKVDL9 and SD4000 had an enhanced ability to invade neurons, survive, and grow intracellularly within the cytosol compared to reference strains such as EGDe. To test this, we grew *L. monocytogenes* to either log phase or stationary phase and then performed gentamicin protection invasion assays in two different neuronal cell lines. Very few gentamicin-resistant *L. monocytogenes* were recovered from N2a murine neuroblastoma cells, with an average invasion rate of only 0.1-0.2% of the inoculum regardless of whether log phase (Fig. 3A) or stationary phase (Fig. 3B) bacteria were used. In F11 dorsal root ganglion-like cells, log phase bacteria invaded more efficiently than stationary phase *Listeria*, but the overall invasion rate was still low and there was no significant difference between the neurotropic and non-neurotropic strains. Since these cell lines are maintained in an undifferentiated state, it was possible that they were lacking expression of a key receptor for bacterial invasion. Thus, we cultured N2a cells in low serum media containing retinoic acid to differentiate them into cells with neurite projections (Fig. 3C), a procedure that makes the cells more closely resemble neurons (23). However, this did not improve invasion for any of the bacterial strains tested (Fig. 3D).

**FIG 3.**
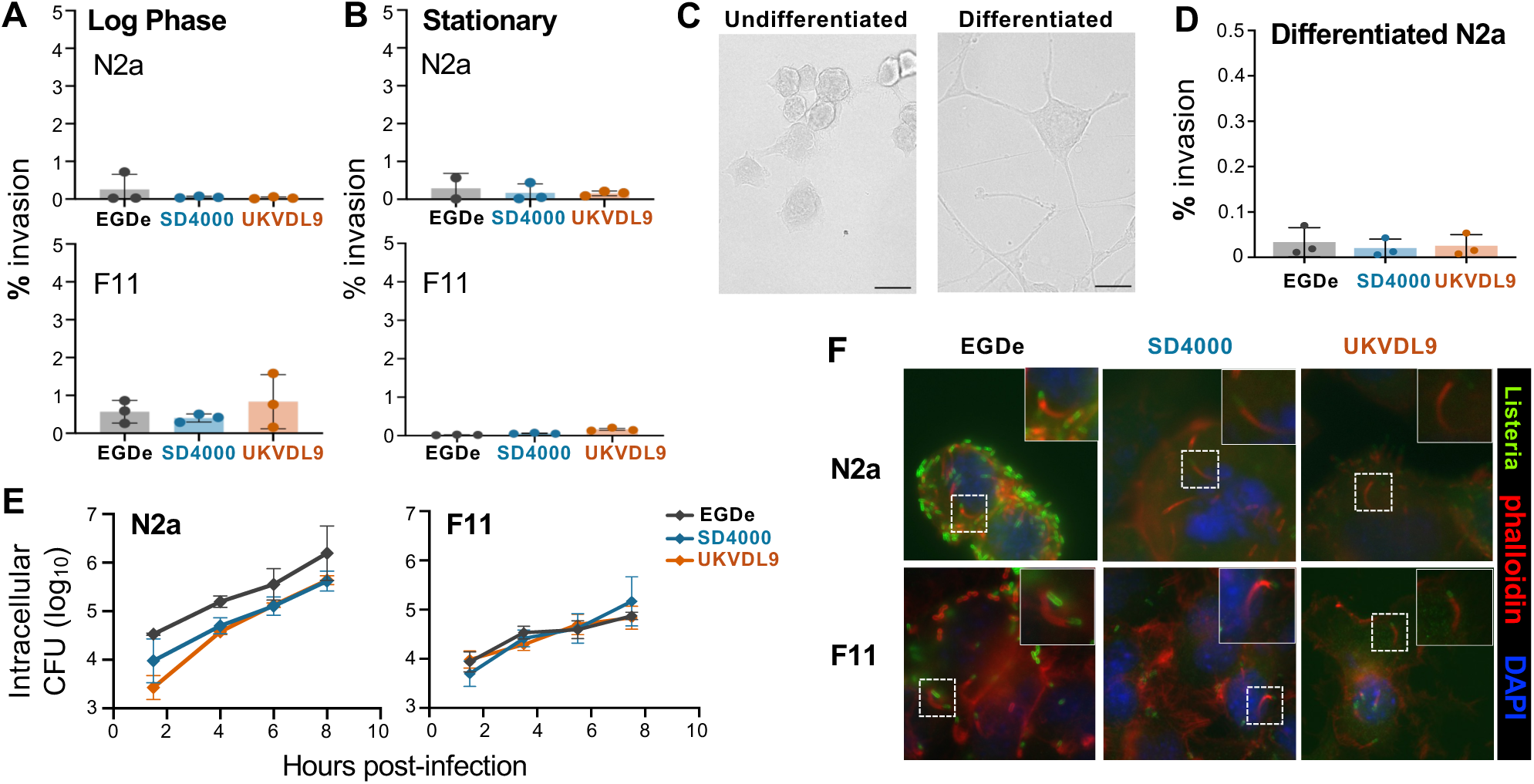
*L. monocytogenes* (*Lm*) invades neurons inefficiently, but the cells can support intracellular growth and actin-based motility. The indicated *Lm* strains were grown to mid-log **(A)** or stationary **(B)** phase and then used to infect N2a or F11 neuronal cell lines at MOI=10. At 1.5 hpi, gentamicin-resistant CFU were calculated as a percentage of the inoculum (% invasion) used to infect the cells; bars indicate mean values (+/− SD) for triplicate samples for one of three independent experiments in each graph. Data were analyzed by one way ANOVA. **(C)** Representative images for N2a differentiated into cells with neurite projections by growth in low serum media containing retinoic acid. Scale bar, 20 µm. **(D)** Invasion assay using stationary phase bacteria in differentiated N2a cells. Symbols represent mean values (+/− SD) for quintuplicate samples from three separate experiments; bars indicate mean values. **(E)** Intracellular growth assay for the indicated *Lm* strains in neuronal cell lines. Symbols represent mean values of triplicate wells from one of a total of three separate experiments performed. Data were analyzed by two-way mixed ANOVA and no differences in the slopes were noted. **(F)** Phalloidin staining was performed at either 4 h (F11 cells) or 6 h (N2a cells) post-infection; dashed lines indicate the zoomed in region shown in the inset.

Although *L. monocytogenes* did not efficiently invade neurons, both the N2a cells and the F11 cells were able to serve as intracellular growth niches for the few bacteria that were able to invade. The number of gentamicin-resistant bacteria increased by nearly two logs over 8 hours in N2a cells (Fig. 3E), a growth rate similar to that observed in macrophages. Growth was more moderate in F11 cells, but still represented a 10-fold increase within this time period. No significant differences were observed in the growth rates of strains EGDe, UKVDL9 and SD4000 in either cell type (Fig 3E). Cytosolic localization of the bacteria in both N2a cells and F11 cells was confirmed by visualizing actin tails at 4-6 hours post-infection (Fig. 3F). Together, these data indicated that *L. monocytogenes* were unable to mediate efficient invasion of neurons, but the few bacteria that were internalized were able to replicate exponentially in the cytosol. Thus, the selective ability of UKVDL9 and SD4000 to disseminate to the brain from the gut was not due to enhanced invasion of neurons as compared to the reference strain EGDe.

### Neurotropic strains display enhanced invasion of enteric glial cells

Given that neurons are typically found surrounded by glial cells, we next considered the possibility that neurotropic *L. monocytogenes* would exhibit an enhanced ability to first invade glial cells and then use ActA-mediated cell-to-cell spread to access the cytosol of neurons. In the gut, a specialized population of cells known as enteric glial cells (EGC) support intestinal neurons. EGC more closely resemble astrocytes than microglia and are characterized by expression of glial fibrillary acidic protein (GFAP) (24). To assess the ability of *L. monocytogenes* to invade EGC, we used a transformed cell line (EGC/PK060399egfr) derived from a rat jejunum (Fig. 4A).

**FIG 4.**
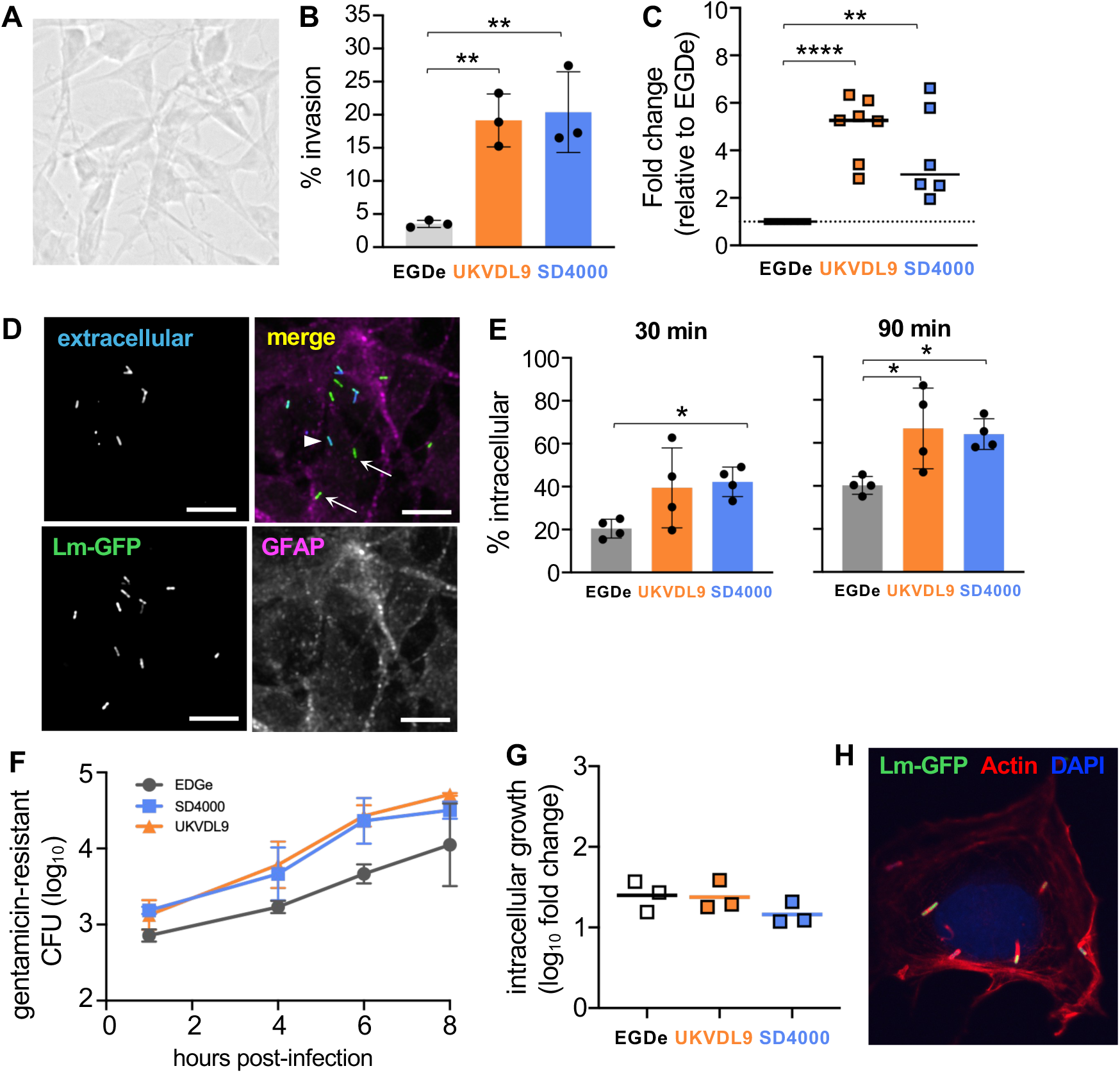
Neurotropic strains UKVDL9 and SD4000 invade enteric glial cells more efficiently than reference strain EGDe. EGC/PK060399egfr cells were infected with the indicated *L. monocytogenes* strains at MOI=5. **(A)** Phase contrast image of uninfected EGC/PK060399egfr cells. **(B)** The percentage of the inoculum that was gentamicin-resistant 1 h post-infection was determined. Representative invasion assay performed with technical triplicates is shown; bars indicate mean values +/− SD. **(C)** Pooled data for n=7 invasion assays is displayed; symbols indicate the mean fold increase in invasion for technical replicates relative to the invasion observed for EGDe in each assay. **(D)** Representative composite and single channel monochromatic images of in-out staining performed on EGC/PK060399egfr cells infected at MOI =0.1 for 90 min. Extracellular bacteria appear blue (arrowhead) and intracellular bacteria appear green (arrow). Sacle bars, 10 μm. **(E)** Quantification of the in-out staining at 30 min and 90 min post-infection for one of two independent experiments. **(F)** Intracellular growth in EGC/PK060399egfr cells was assessed by measuring gentamicin-resistant bacteria at each time point. A representative assay performed with technical quadruplicates in shown. **(G)** Compiled data for three separate intracellular growth assays is shown as the fold increase in total gentamicin-resistant CFU over over 8 hours. **(H)** Representative image showing cytosolic *L. monocytogenes* with actin tails 9 hpi in EGC/PK060399egfr cells. Data in panels B, C, E, and G were analyzed by one-way ANOVA with Dunnett’s multiple comparisons test.

Both UKVDL9 and SD4000 invaded the EGC cell line significantly better than reference strain EGDe within 1 hour post-infection (Fig. 4B). On average, the neurotropic bacteria were internalized 3 to 5-fold more than the non-neurotropic strain (Fig. 4C). We also confirmed the results of the gentamicin protection assays with a microscopic approach to distinguish intracellular and extracellular bacteria (Fig. 4D). By 30 minutes post-infection, xxx was seen and at 90 minutes Fig. 4E – waiting on River’s analysis of the repeat experiment

All three bacterial strains tested were able to replicate exponentially in the cytosol of the EGC cell line (Fig. 4F). Regardless of the strain used, the cells supported a 20-fold increase in CFU over an 8 hour period (Fig. 4G), a *L. monocytogenes* growth rate comparable to that seen in epithelial cells. At 9 hours post-infection, *L. monocytogenes* in the EGC cell line were observed with actin tails (Fig. 4H). Together, these results indicated that neurotropic strains UKVDL9 and SD4000 had an enhanced ability to invade the EGC cell line, and that they were actively replicating in the cytosol and were equipped for cell-to-cell spread.

To ensure that the results obtained above were not an artifact from using a transformed cell line, we next attempted to isolate primary enteric glial cells from murine small intestines. Briefly, the muscularis was physically removed from the duodenum and jejunum, and then the cells were chelated with EDTA to remove intestinal epithelial cells and digested with collagenase (Fig. 5A). Non adherent cells were washed away 24 hours later and the selective growth of EGC was promoted by culturing the cells for several days in low serum media containing specific neural supplements. Within a few days, clumps of cells with the morphological characteristics of EGC appeared (Fig. 5B) and the cells became confluent with continued in vitro culture. The primary cells expressed similar levels of GFAP as the PK060399egfr cell line (Fig. 5C) and on average, were greater than 90% pure (Fig. 5D).

**Fig. 5.**
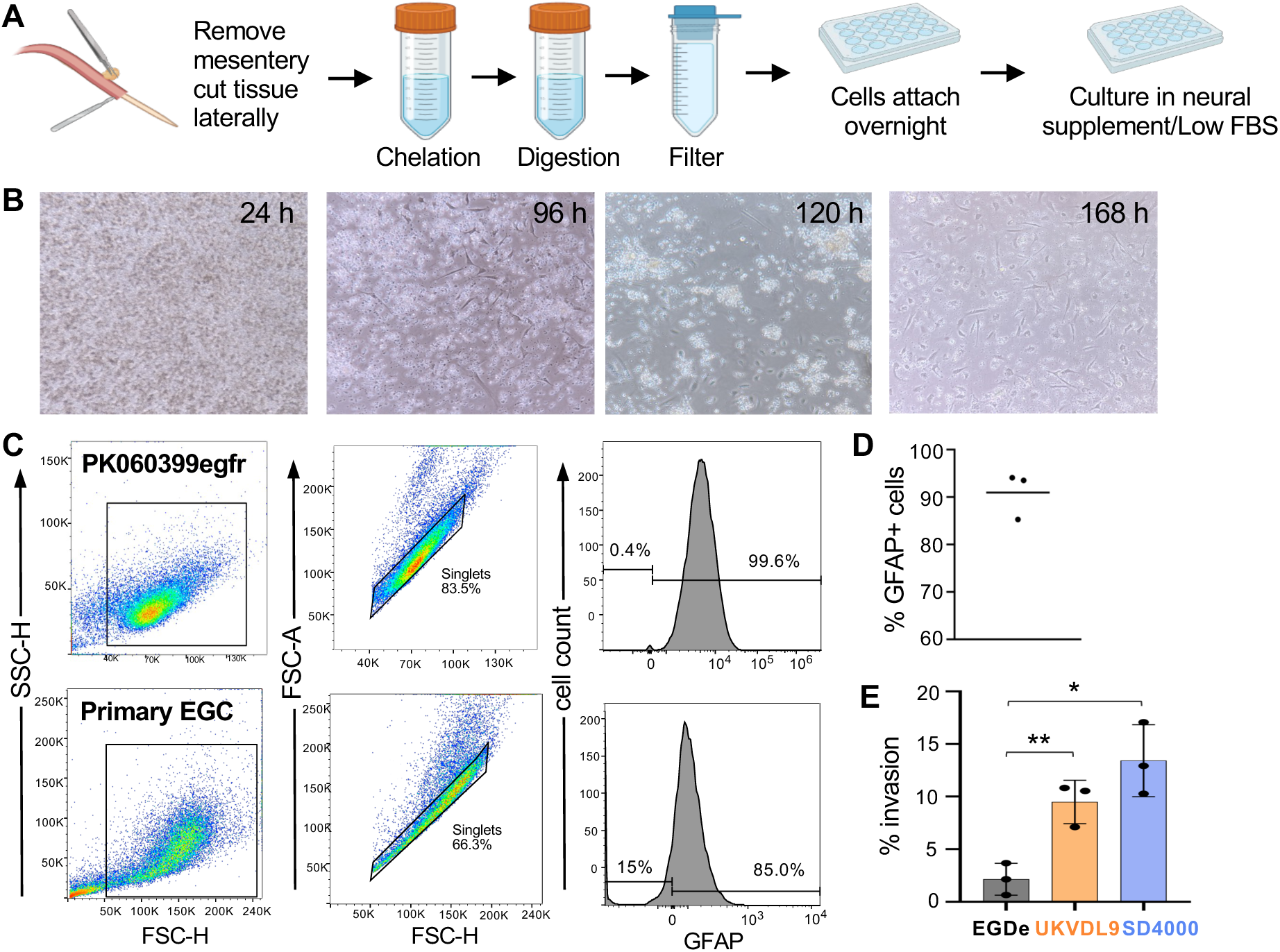
Neurotropic strains UKVDL9 and SD4000 invade primary enteric glial cells (EGC) more efficiently than reference strain EGDe. **(A)** Outline of the procedure to isolate EGC from murine small intestines. **(B)** Representative images of primary EGC during the various stages of the in vitro culture process. **(C)** Representative flow cytometry gating strategy for identification of glial fibrillary acidic protein (GFAP)-positive EGC. The top row shows results for the rat cell line EGC/PK060399egfr and the bottom row shows primary murine EGC. **(D)** Average purity of the primary EGC used for invasion assays. (E) Primary EGC were infected with the indicated *L. monocytogenes* strains at MOI=1 and the number of gentamicin-resistant bacteria was determined 1 h later. Symbols represent mean values for two technical replicates from three independent experiments; bars indicate mean values +/− SD. Data were analyzed by one-way ANOVA with Dunnett’s multiple comparisons test.

Confluent wells of primary EGC were infected with *L. monocytogenes* and the percent of the inoculum that was internalized 1 hour post-infection was determined. As shown in Fig. 5E, both UKVDL9 and SD4000 invaded the primary EGC more efficiently than reference strain EGDe. Over three separate experiments using different batches of primary enteric glial cells, we found that the 5-to-6-fold more neurotropic *L. monocytogenes* strains were internalized than the non-neurotropic strains. Thus, the result seen with the transformed EGC cell line was confirmed in primary cells, validating the use of the EGC/PK060399egfr cell line for further studies.

### Neurotropic *Listeria* use actin-based motility to spread from enteric glial cells to neurons

To determine if *L. monocytogenes* were capable of spreading from enteric glial cells into neurons, we developed an *in vitro* cell-to-cell spread assay in which the EGC cell line was infected with *L. monocytogenes* and then overlaid onto differentiated N2a cells (Figure 6A). A ratio of 4:1 enteric glial cells to neurons was used to maximize cell adjacency and maintain a physiologically relevant distribution of cells (25). The co-cultured cells were maintained in gentamicin (40 µg/mL) to ensure that any extracellular bacteria would not be able to replicate, and thus, would not be able to form actin tails. Neurons were visualized by staining with β-3-tubulin. The enteric glial cells were not stained with any specific marker but the phalloidin used to visualize actin tails associated with GFP-expressing *L. monocytogenes* also stained the diffuse actin network making the cell boundaries of the β-3-tubulin-negative enteric glial cells easy to determine. Extracellular bacteria were distinguished by co-labeling with endogenous GFP and an anti-*Listeria* antibody. Images were collected from various timepoints from 6 to 24 hours post-infection across four separate experiments. Analysis of the images revealed that there was little phenotypic difference in images collected between 6 and 12 hours post-infection, likely due to the fact that individual cell-to-cell spread events happened asynchronously within infectious foci. Therefore, we grouped any image taken from t=6-12 hours as early cell-to-cell spread timepoints. Likewise, the images collected from 18 to 24 hours post-infection were classified as late cell-to-cell spread events (Figure 6A).

**Fig 6.**
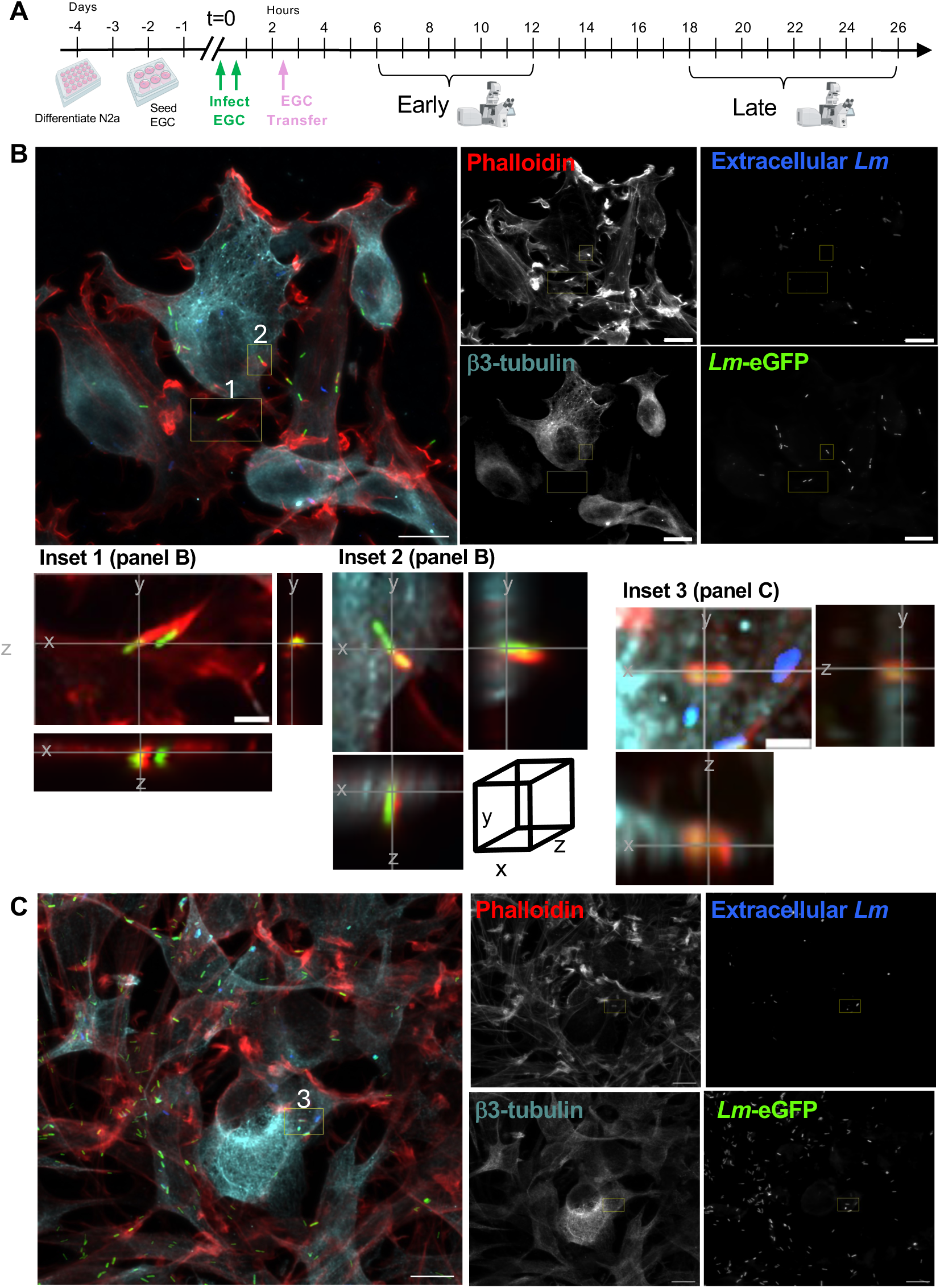
*L. monocytogenes (Lm)* demonstrate actin-based motility in infected EGC and were occasionally found inside N2a cells 4-8 hours after transfer. **(A)** Timeline for the cell-to-cell spread assay in which infected EGC (EGC/PK060399egfr cell line) were overlayed onto differentiated neurons (N2a cells) in a 4:1 ratio in media containing 40 μg/ml gentamicin. The EGC were infected with eGFP-expressing *Lm* UKVDL9 (green) at a MOI of 0.25. N2a cells were identified by β_3_-tubulin staining (cyan); actin was visualized with phalloidin (red); and extracellular bacteria were identified by staining non-permeabilized cells with anti-*Lm* antibody (blue). Four independent experiments were performed with multiple time points imaged in each experiment. Representative images from one of four independent experiments depicting the early time period (4-8 hours after cell transfer) are shown in this figure and the late time period (16-24 h after cell transfer) in Fig. 7. **(B)** A merged image from t=10 h and the corresponding single channels are shown. Orthogonal projections (10μm) of the insets marked (1) and (2) are shown below the merged image. (**C**) A merged image and monochromatic channels from t=6 h; orthogonal projections of the inset marked (3) are shown above. Scale bars in composite and single channel images, 10μm; in orthogonal projections, 2μm.

The enteric glial cells were harvested at t=2.5 hours when the bacteria contained within them would be localized to the cytosol and surrounded by actin clouds due to the uniform distribution of ActA on the surface, but not yet replicating exponentially and forming actin tails (Fig. 4). The early phase time points (6-12 hours post-infection but only 3.5 to 9.5 hours post co-culture) were chosen with the expectation that the majority of EGC would have multiple motile bacteria in the cytosol. Thus, we expected the early time points to show the initial movement of *L. monocytogenes* from the enteric glial cell into the neurons. Indeed, enteric glial cells that contained many bacteria with actin clouds were readily observed at early time points (Fig. 6B, inset 1). It was common to find *L. monocytogenes* with actin tails that were approaching the bounds of an N2a cell (Figure 6B, inset 2). Less commonly, *L. monocytogenes* were visualized inside the neurons, surrounded by an actin cloud (Fig. 6C, inset 3). These observations indicated that spread from EGC to N2a cells was beginning to happen during the early stage of the assay, but suggested there had not been enough time for the bacteria to begin replicating in the cytosol of the neurons.

At the late time points (t=18-26 hours post-infection and 15.5-23.5 hours post-culture), we expected to find robust intracellular replication of *L. monocytogenes* and significant bacterial spread both from EGC to neurons and between EGC. This prediction was based on previously published cell-to-cell spread assays using transformed cell lines of epithelial origin (26) or spread from macrophages to rat neurons (27). As demonstrated by the representative image shown in Fig 7A, the N2a cells contained numerous *L. monocytogenes* with long actin tails in the soma. Longitudinal views (Fig 7A, inset 1) showed that these bacteria were completely contained within the bounds of the neuron (Fig. 7A). The large number of intracellular bacteria observed in the N2a cells at the later timepoints suggested that the cell-to-cell spread events happened several hours earlier and that there had been time for both replication of *L. monocytogenes* to occur in the cytosol and subsequent polarization of ActA on the cell surface. These conclusions were also supported by the intracellular growth assays shown in Fig. 3, which showed a 100-fold increase in bacterial growth over eight hours in N2a cells.

**Fig 7.**
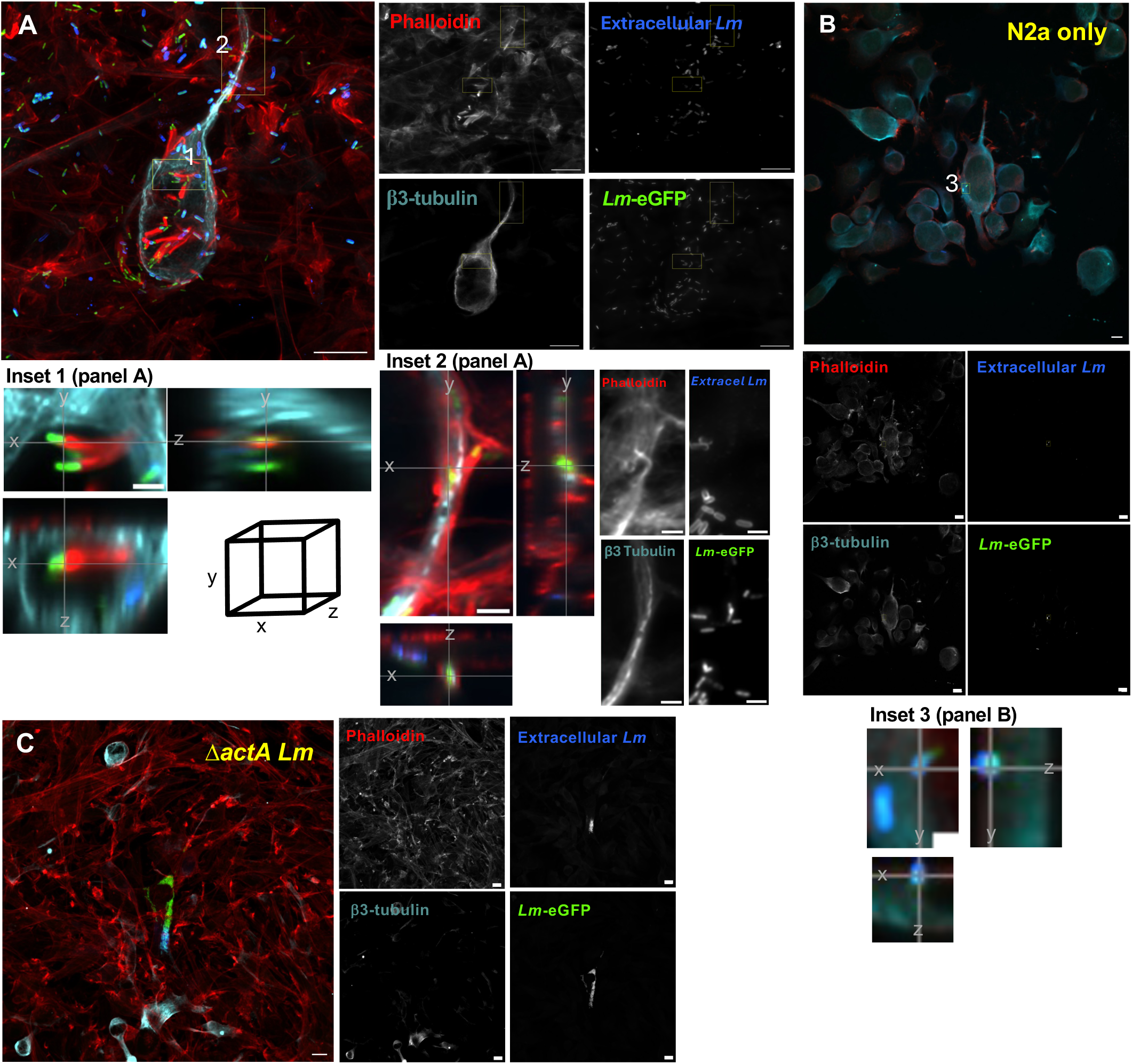
*L. monocytogenes* uses ActA to spread from enteric glial cells to neurons. A cell-to-cell spread assay was performed as described in Fig. 6A. Representative images from one of four independent experiments are shown. The EGC were infected with eGFP-expressing *Lm* UKVDL9 (green); N2a cells were identified by β_3_-tubulin staining (cyan); actin was visualized with phalloidin (red); and extracellular bacteria were identified by staining non-permeabilized cells with anti-*Lm* antibody (blue). **(A)** A merged image representative of the late time period and single channel monochromatic channels from t=21 h. Orthogonal projections of insets (1) and (2) are shown below. **(B)** Composite image and single channel monochromatic channels of control coverslips containing N2a cells only at t=6 h; orthogonal projections of inset (3) are shown below. **(C)** Composite and single channel images from control coverslips infected with eGFP-expressing *ΔactA Lm* fixed at t=10 h. Scale bars in composite and single channel images, 10μm; in orthogonal projections, 2μm.

We also observed a large number of *L. monocytogenes* localized within the bounds of a neurite or forming protrusions from the neurite (Fig. 7A, inset 2; Supplemental Movie, Fig. S1A). These bacteria were trafficking both toward and away from the soma. Interestingly, we found many *L. monocytogenes* associated with actin tails or actin clouds near the distal terminus of N2a neurites, particularly when that neurite was located near or within the bounds of an adjacent enteric glial cell. These observations suggested that the distal end of an axonal projection may serve as a potential site for cell-to-cell transfer from EGC or sites of secondary reinfection from N2a back to EGC (Fig. 7A, inset 1 and Figs. S1A and S1B).

In the late stages of co-culture, it was common to see many attached extracellular bacteria, as well as some intracellular bacteria lacking actin tails in the EGC surrounding the neurons. These observations would be consistent with a heavily infected cell having lysed and the resulting extracellular bacteria being briefly exposed to gentamicin before subsequently being taken up by EGC where they were unable to replicate intracellularly due to an inhibition of protein synthesis. To determine if the gentamicin pressure was sufficient to prevent intracellular replication of any *L. monocytogenes* that were exposed to the media, we infected differentiated N2a monolayers in the presence of gentamicin (40 µg/mL) and collected coverslips during both the early and late time periods. Only extracellular bacteria were found associated with N2a cells 10 hours post-infection (Fig. 7B). By 17 hours post-infection, the extracellular *L. monocytogenes* were almost completely eliminated (Fig. S1C).

To determine whether the spread from enteric glial cells to neurons was dependent on actin-based motility, we infected the EGC cell line with a Δ*actA* variant of *L. monocytogenes* UKVDL9. In these co-cultures, the bacteria lacking ActA remained inside enteric glial cells and were unable to spread to N2a cells at either early (Figure 7C) or late (data not shown) time points. Together, these results clearly demonstrate the capacity of *L. monocytogenes* to efficiently invade enteric glial cells, replicate intracellularly, and then spread to adjacent neurons, a dissemination mechanism that positions the bacteria to then use actin-based motility to move along the length of an axon to the brainstem.

## DISCUSSION

A subset of *L. monocytogenes* strains can use actin-based motility to disseminate from the gut directly to the brainstem to cause rhombencephalitis in a disease process that is thought to be linked more closely to the virulence of the bacteria than the immune status of the host. In this report, we showed that two of these *L. monocytogenes* strains did not efficiently invade neurons directly. Instead, the neurotropic strains displayed a preferential ability to invade the enteric glial cells that surround intestinal neurons. Using an *in vitro* co-culture system, we demonstrated that *L. monocytogenes* could readily spread from the cytosol of an enteric glial cell into the cytosol of a terminally differentiated neuron. The results presented here provide further evidence that the foodborne mouse of model of listeriosis can be used to differentiate the distinct mechanisms of dissemination that lead to either diffuse meningitis or localized rhombencephalitis infections and provide initial insights into how the neurotropic strains of *L. monocytogenes* access the nervous system.

Intravenous transmission of any *L. monocytogenes* strain that can achieve a titer of 100-1000 CFU/mL in the blood can directly invade the blood-brain barrier (7), and we posit that either some degree of immune compromise in the host or use of an overwhelming inoculum is needed for this to occur. In contrast, a subset of neurotropic strains of *L. monocytogenes* encode virulence factors that allow for invasion of cranial nerves in the gut (28, 29) and potentially an altered form of actin-based motility (30) to promote axonal migration the length of the nerve to the brainstem. *L. monocytogenes* strains have been grouped into four phylogenomic lineages and the two neurotropic strains used in this study both belong to phylogenomic lineage III (4). The majority of isolates sequenced to date are classified as either lineage I or lineage II (2); therefore, relatively less is known about the pathogenic potential of lineage III and lineage IV strains. Lineage III was initially described as being in higher proportions in veterinary isolates than in human clinical samples (31) and has been linked to severe outbreak of disease in cattle (32). Rhombencephalitis is a common presentation of listeriosis in ruminants so it is possible that cows and sheep serve as a reservoir for neurotropic strains that can cause brainstem infections in otherwise healthy humans (10). However, multiple studies that analyzed the sequence type of rhombencephalitis isolates from ruminants found that the majority were classified as lineage I (33–35), following the same pattern observed for other manifestations of listeriosis in both humans and ruminants. Thus, the ability to invade enteric glial cells, spread to neurons, and then ascend to the brainstem is likely to be caused by particular virulence genes present in the accessory genome of neurotropic strains rather than due to an association with a particular lineage.

The most likely route of spread from the intestines to the brainstem is the vagus, a cranial nerve with about 40,000 axons (in humans) that innervate the upper GI tract (36). Vagal afferent endings are distributed in all layers of gut tissue (37): lamina propria, submucosal plexus, and myenteric plexus and we showed here that *L. monocytogenes* were found in each of these layers of the ileum following foodborne transmission. Ruminants are thought to acquire *L. monocytogenes* through consumption of contaminated silage and since they regurgitate partially digested food and chew it again, cranial nerves that innervate the upper gastrointestinal tract including the oral cavity may also be involved. We did attempt to harvest bacteria in the process of traversing the vagus nerve, but this approach is technically challenging and the lack of recovery of bacteria in many mice cannot be interpreted as evidence against the use of this pathway in the mouse model. It takes only a small number of founder bacteria to cause brain infection with significant symptoms and there is a period of four to five days during which the bacteria could be present in the vagus nerve prior to colonizing the brain. We still do not know if this delay is due to the time needed for the bacteria to invade enteric glial cells and spread into neurons, or if it because the axonal migration along the length of the cranial nerve requires an extended time period.

To the best of our knowledge, this is the first time that enteric glial cells have been directly implicated in the pathogenesis of a bacterial infection in the gut. Previous studies that examined enteric glial cells during infection focused primarily on cytopathic responses that occurred where EGC were not the primary target for pathogen replication. For example, Trevizan et al. showed that oral infection of Wistar rats with *Toxoplasma gondii* resulted in an increased ratio of enteric glial cells to neurons in the duodenum due to a loss of neurons in the submucosal plexus(38). Likewise, Janova et al. found that murine enteric glial cells were not a direct target of West Nile virus, but they contributed to the inflammatory response during intestinal infection (39). When primary human enteric glial cells were co-cultured with Caco-2 cells, fewer *Shigella flexneri* infectious foci in the epithelial cells and less epithelial barrier damage were observed (40). In that study, it was suggested that the EGC were not directly infected because they down-regulated Cdc42 which is required for *S. flexneri* invasion of epithelial cells. In contrast, the results of this study suggest that the ability to invade enteric glial cells may be critical for neurotropic strains of *L. monocytogenes* to gain access to cranial nerves in the gut. We propose that direct infection of neurons is not likely to occur frequently *in vivo* both because of the low invasion rate and the fact that they are enveloped by the supporting enteric glial cells.

The rat enteric glial cell line used in this report was previously used to investigate *Clostridioides difficile* toxins TcdA and TcdB. In those studies, the *C. difficile* toxins caused rapid cytopathic effects and apoptosis of the cells within 24 hours of treatment (41–43). Although TcdA and TcdB were already known to affect epithelial cells and neurons, the authors noted the important role of enteric glial cells in gut homeostasis and suggested that their susceptibility to the toxins could play a role in the lingering intestinal dysfunction observed in patients following *C. difficile* infection. In that study, the cells were used until passage 20; however, we found that the enhanced invasion phenotype for neurotropic *L. monocytogenes* in this cell line became less reliable after passage 12.

Enteric glial cells are most similar in structure to astrocytes, the star-shaped glia found only in the brain or spinal cord, but they also have unique features not found in any of the three known glial cell types of the CNS (44, 45). For example, enteric glial cells are important for both normal development of the intestines and maintaining gut homeostasis. Secreted factors produced by enteric glial cells influence both epithelial cell proliferation and barrier function (46), and the role of the gut microbiota in regulating these processes is only beginning to be established (47). In addition, EGC respond to either injury or inflammation by secreting cytokines that can influence the immune response. Previous work showed that primary human EGC exposed to *Shigella* or enteroinvasive *E. coli* upregulated nitric oxide release in a TLR-dependent manner (48). Although there had been some suggestion that EGC may serve as unconventional antigen presenting cells, Brown et al. recently showed that the cells presented antigen only on MHC-I and not MHC-II which was expressed at only a low level on EGC during *Toxoplasma gondii* infection (49). Thus, the emerging consensus is that enteric glial cells are ideally positioned in the gut to alter cellular responses in both homeostasis and during inflammation, and future studies will be needed to elucidate their exact role during infection with *L. monocytogenes* and other intestinal pathogens.

## MATERIALS AND METHODS

### Bacterial strains and culture conditions

*L. monocytogenes* strains EGDe, UKVDL9, SD4000 were described previously (4, 11). *L. monocytogenes* strains PF49 and PF12 were generously provided by Jose Vazquez-Boland (University of Edinburgh). An *actA* deletion mutant in the UKVDL9 strain background was generated by allelic exchange using the pKSV7 vector and the resultant strain (*Lm* SD9012) was confirmed by whole genome sequencing (SeqCoast). Variants that constitutively expressed eGFP *Lm* under the control of the P_help_ promoter were generated using pGJ-cGFP (50), resulting in strains SD9900 (UKVDL9::eGFP) and SD9912 (UKVDL9 Δ*actA*::eGFP). All strains were grown in Brain Heart Infusion media (Difco). Frozen aliquots of *L. monocytogenes* were thawed at room temperature and then allowed to recover in BHI broth for 1.5 h standing at 30°C (for *in vivo* infection of mice) or 37°C (for *in vitro* infection of cells) (51).

### qRT-PCR

Bacteria were grown shaking in BHI media at 37°C until late logarithmic phase (OD_600_ = 0.8) and then pelleted and suspended in TRIzol (Invitrogen). RNA was extracted as directed and quantified using a Nanodrop instrument. RNA was pre-treated with ezDNAse (Invitrogen) and cDNA was generated using random primers in the SuperScript IV First Strand Synthesis system (Invitrogen). A control reaction lacking reverse transcriptase was performed for each sample and all cDNAs were treated with RNAseZ (Invitrogen). For qPCR, cDNA was added to KiCqStart SYBR Green qPCR ReadyMix (Sigma-Aldrich) and forward and reverse primers (10 μM) were added in triplicate to a MicroAMP 96-well plate (Applied Biosystems). Amplification was carried out by CFX Opus 96 Real-Time PCR System (BioRad). Oligonucleotide primers (IDT DNA, Inc.) used were: 5’-AAGCACAACCCAAGAAGGAA-3’ (*inlB* forward); 5’-CGGTGATAGTCTCCGCTTGT-3’ (*inlB* reverse); 5’-TTAGCTAGTTGGTAGGGT-3’ (16S forward); 5’-AATCCGGACAACGCTTGC-3’ (16S reverse). Expression of *inlB* was normalized to 16S rRNA expression for each sample using the 2^-ΔΔCt^ method and fold change relative to the mean value for EGDe was reported.

### Mouse models of infection

BALBc/ByJ mice (stock #001026) were purchased from The Jackson Laboratory and a small breeder colony was maintained in SPF conditions at the University of Kentucky. Four week old females were transferred to a room with a dark cycle from 9 AM – 7 PM and given at two weeks to acclimate to the light cycle prior to infection. For i.v. inoculation, bacteria suspended in a total volume of 200 µl of Dulbecco’s phosphate buffered saline (PBS; Invitrogen #14190) were injected into the tail veins of mice. For foodborne infection, mice were housed on raised wire flooring, fasted for 18 hours to prevent coprophagy, and then fed a small piece of bread contaminated with *Listeria* as described (52). All procedures were approved by the University of Kentucky Institutional Animal Care and Use Committee.

### Processing of tissues for CFU determination

Spleens, livers, and brains were harvested aseptically and homogenized in sterile water for 30 sec using a PowerGen 1000 homogenizer at 60% power. Serial dilutions were prepared in sterile water, plated on BHI agar, and incubated overnight at 37°C. Colons were flushed with 8 mL of sterile PBS, opened longitudinally with a scalpel, cut into small fragments, and then homogenized for 60 sec at 80% power. Serial dilutions were prepared in sterile water, plated on BHI L+G agar (53), and incubated for two days at 37°C. Anterior and/or posterior segments of the vagus nerve (5-10 mm) extending from the vagal trunk and extending along the esophagus were dissected and placed in 1 mL of 0.2% NP-40 in a 2 mL Dounce homogenizer (Bellco). The homogenized tissue was transferred to a centrifuge tube along with 8 mL of sterile water that was used to wash out the homogenizer and then the bacteria were pelleted by centrifugation at 26,953 *x g* for 15 min. Serial dilutions were prepared in sterile water, plated on both BHI agar and CHROMagar^TM^ *Listeria*, and incubated overnight at 37°C. For all samples, *L. monocytogenes* burdens were expressed as total CFU per tissue.

### Co-localization of *Listeria* with neurons

Small intestines were aseptically harvested from infected mice and the terminal third was designated as the ileum. Ileums were flushed with 10 mL of modified Bouin’s fixative (50% ethanol 5% acetic acid in water), cut longitudinally, and then wrapped around a sterile toothpick starting with the proximal end. The rolled tissue was fixed with paraformaldehyde, dehydrated using glycine and sucrose, embedded in OCT, and stored at −70°C as described previously (54). Thin sections (10 μm) were washed 3X with a solution consisting of 1% BSA, 0.2% Triton-X-100 in PBS, and then blocked by adding 10% normal goat serum, 3% bovine albumin, 0.1% Triton-X, and 0.05% Tween-20 in PBS. Sections were stained with rabbit *Listeria* O antiserum polyclonal (1:2000, BD Difco) overnight at 4°C, AF594-conjugated goat anti-rabbit secondary antibody (1:200; Invitrogen) for 2 h, and 1:200 AF-488 conjugated β_3_-tubulin antibody (1:100; polyclonal, Millipore Sigma) for 2 h. After three final washes with the wash buffer, ProLong Diamond Antifade Mountant with DAPI (Invitrogen) was applied. Blinded slides were imaged using 60X and 100X objectives on an EVOS M5000 Imaging System.

### Cell culture

Rat enteric glial cells (EGC/PK060399egrf) were purchased from ATCC (CRL-2690) and used until passage 12 by culturing in DMEM (Life Technologies Cat. #11960) with 2 mM L-glutamine, 1 mM sodium pyruvate and 10% fetal bovine serum (FBS; Gemini). N2a cells were provided by John Gensel and cultured in media consisting of 40% DMEM with high glucose and pyruvate (Life Technologies cat. # 10313), 50% Opti-MEM (Life Technologies cat. # 31985), and 10% FBS. To differentiate the cells, N2a were seeded in 24-well culture plates and allowed to grow for 24 hours before changing the media to differentiation medium which consisted of 45% DMEM, 54% Opti-MEM, 1% FBS and 10 µM retinoic acid (Sigma). The cells were allowed to differentiate for 72 hours prior to infection, replacing the media every 24 hours. F11 cells were provided by John Gensel and cultured in DMEM with high glucose and pyruvate (Life Technologies cat. # 10313) and 10% FBS. All cells were routinely passaged in media containing both penicillin (10 U/mL) and streptomycin (10 µg/mL) (Life Technologies cat. #15140122), and the cells were washed once and media lacking antibiotics was added at least several hours prior to infection.

### Intracellular invasion and growth assays

Cells were seeded in antibiotic-free media in a 24-well tissue culture treated plate (Corning) at 37°C with 5% CO_2_ and cultured overnight to achieve 80-90% confluency. Stationary phase bacteria were washed once with sterile PBS and diluted to achieve the desired multiplicity of infection (MOI) for an inoculum in 50 µl per well. Plates were centrifuged at 300 x *g* for 5 minutes to synchronize the infection. At either 1 h post-infection (EGC cells) or 1.5 h post-infection (N2a and F11 cells) media containing 15 µg/mL of gentamicin was added for 20 min at 37°C/5% CO_2_ to kill extracellular bacteria and then the cells were washed with pre-warmed PBS. To lyse the cells, 1 mL of cold sterile H_2_O was added, the plate was incubated for 15 min at 4°C, followed by vigorous pipetting up and down. Serial dilutions of the cell lysates were plated on BHI agar and the percent inoculum was determined by dividing the total number of intracellular bacteria recovered by the number of bacteria in the inoculum (which was determined precisely by preparing serial dilutions and plating). Intracellular growth over time was reported as the total number of gentamicin-resistant bacteria recovered at each time point.

### In/out staining to identify intracellular bacteria

EGC were cultured in antibiotic-free media on 12 mm round glass coverslips in 24-well plates at 37°C with 5% CO_2_ and infected with *L. monocytogenes*. Coverslips were washed twice with 1 ml of prewarmed PBS and fixed in 4% buffered paraformaldehyde for 15-20 minutes. Cells were blocked in 300 µl of antibody buffer (TBS + BSA 1% w/v) for 30 min at room temperature (RT). After 30 min, cells were washed 8X with 300 μl of antibody buffer. Listeria O Antiserum (Polyclonal Rabbit IgG, Invitrogen) diluted 1:100 in antibody buffer was added to each well and incubated at RT for 45-60 min. Cells were washed 8X with 300 μl of antibody buffer and then 200 µl secondary antibody (AF405-conjugated goat anti-rabbit diluted 1:100; ThermoFisher) was added and incubated at RT in the dark for 30-60 min. Cells were washed 8X with 300 μl of antibody buffer and then permeabilized with 0.1% Triton X-100 for 30 min at RT in the dark. After 30 minutes 1:40 ul of A GFAP specific antibody (eFlour-660-conjugated, clone GA5, ThermoFisher or BV421-conjugated, clone: 2E1.E9, BioLegend) diluted 1:40 in antibody buffer containing detergent was added for 40 min at RT in the dark and then the coverslips were washed 8X first with TBS containing 0.1% Triton X-100 then 8x with TBS alone. After drying, the coverslips were mounted with ProLong Diamond anti-fade and images were collected on a Nikon AXR Inverted Confocal Microscope and analyzed in Nikon Element version 6.10.0 and ImageJ version 1.54p

### Visualization of actin tails

Cells were cultured in antibiotic-free media on 12 mm round glass coverslips in 24-well plates at 37°C with 5% CO_2_ and infected with *L. monocytogenes* at a MOI of 10. Plates were centrifuged at 300 x *g* for 5 minutes to synchronize the infection. At 1.5 h post-infection, gentamicin was added at a final concentration of 30 µg/ml and the cells were further incubated at 37° C/5% CO2. At the indicated time points, cells were washed once with PBS, fixed with formalin for 15 min at RT, washed 3X with PBS, and then permeabilized with Tris-buffered saline (TBS) containing 0.1% Triton X-100 v/v and 1% BSA w/v (TBS-TX) for 30 min. Cells were stained with rabbit Listeria O Antiserum Poly (1:2000; BD Difco) and Texas Red-phalloidin (1:100; ThermoFisher) in TBS-TX for 30 min in a dark, humid chamber at RT. After washing 8X with TBS-TX, AF488-conjugated anti-rabbit secondary polyclonal antibody (1:200) was added for 45 minutes. Cells were washed 8X with TBS-TX, then washed 8X with TBS only. Once dry, coverslips were mounted on slides with Prolong Diamond Antifade with DAPI (Life Technologies) and visualized with the 60x objective of an EVOS M5000 Imaging System.

### Isolation of primary enteric glial cells

The following protocol was adapted from previously published reports (55, 56) based on pilot optimization studies and personal communications with one set of authors (56). Poly-L-lysine-coated plates were prepared by adding 400 µl (for a 24 well dish) or 1 mL (for a 6 well dish) per well of 0.5 mg/mL solution and incubating either overnight at room temperature or for 1 h at 37°C. Plates were washed twice with sterile distilled H_2_O and allowed to dry either overnight at room temperature or for 2 h at 37°C. Small intestines were harvested from BALBc/ByJ mice (both male and female mice, aged 3-5 months were used) and the proximal first third (duodenum) and middle third (jejunum) were processed separately. Intestinal segments were flushed with 8 mL of cold PBS (Life Tech cat. #14190) and then a wooden dowel presoaked in ice-cold PBS was inserted. Keeping the tissue moistened with cold PBS, the muscularis was removed by making a small nick on the edge and then using a moistened sterile swab to gently roll it the end of the tissue and detach with forceps. Each segment was then cut longitudinally and lifted off the wooden dowel.

Segments were cut into fourths, and all the segments from 2 mice were combined in a 50 mL centrifuge tube containing 30mL of chelation buffer (PBS with 2 mM EDTA, 10 mM HEPES, 1% FBS) and incubated shaking at ∼200 rpm at 37°C in 5% CO_2_ for 20 minutes. Each tube was vigorously shaken up and down four times, then the contents were passed through a 70 µm filter. The supernatant was discarded, and the tissue was washed with 2 mL of room temperature PBS. This chelation procedure was repeated twice more and then the remaining tissue was cut into smaller pieces (< 0.5 cm) and placed into a new tube containing 20 mL of digestion buffer consisting of PBS with 10 mM HEPES, Collagenase type III (1.5 mg/mL; Fisher cat. #7422), and DNAse I (80 µg/mL;Worthington Biochemical cat. #L5002007). The tissues were digested by shaking at ∼200 rpm for 20 min at 37°C in 5% CO_2_, passed through a 40 µm cell strainer and the filtrate was collected and centrifuged at 400 *x g* for 5 minutes at 4°C. The cell pellet was suspended in 3 mL of glial seeding media consisting of DMEM/F12 (Life Tech cat. #11320) with 1X Gem-21 supplement (Gemini Bio cat # 400-160-010) 1X G-5 supplement (Fisher cat #17503012), 1X N-2 supplement (Gemini bio cat. #50-753-3047), 10 mM HEPES, 2 mM GlutaMAx (Gibco cat # 35050-60), 10% FBS 10%, 100 U/ml Penicillin-Streptomycin (Life Tech cat # 15140), and 15 µg/mL gentamicin.

Cells were seeded (500 µl/well) in six wells of poly-L-lysine-coated 24 well dish each containing 500 µL of prewarmed glial seeding media and incubated for 24 h at 37°C with 5% CO_2_. Attached cells were washed once with pre-warmed PBS and then fed 1mL of prewarmed Glial Growth Media, which was similar to the seeding media except for a reduced FBS concentration of only 0.5%. Cells were further incubated at 37°C with 5% CO_2_ for 48 h. After incubation, plates were checked for clumped or grouped cells with glial morphology, and the media was changed to fresh glial growth media every 72 h. Wells were typically 85-95% confluent within 4-11 days post-seeding.

### Flow cytometry

Single cell suspensions were incubated with fluorescently conjugated antibodies specific for CD90.2 (clone 53-2.1) BioLegend), CD45 (clone 30-F11), GFAP (clone 2E1.E9) purchased from BioLegend and CD24 (clone M1/69), beta-3-tubulin (clone 2G10-Gb3), and GFAP (clone GA5) from Invitrogen. Flow cytometry data were acquired using a Cytek Aurora, and analyzed using FlowJo software (Version 10.10.0).

### Cell-to-cell spread assay

N2a cells (passage 6-10) seeded on 12 mm round glass coverslips in 24-well dishes (Corning) and allow to differentiate for 3 days. EGC/PK060399egrf cells (passage 5-7) were seeded into 6-well dishes in media without antibiotics and incubated overnight at 37°C in 5% CO_2_. *L. monocytogenes* strains were prepared as described above and diluted to prepare inocula to infect the EGC cells at a MOI of 0.25 twice (at t=0 and t=30 min) to better distribute the bacteria amongst the cells. At t=1.5 h, CLR-2690 cells were washed once with pre-warmed PBS, and suspended in 1 ml of CRL-2690 media containing gentamicin (40 μg/ml) for 20 min at 37°C with 5% CO_2_ to kill extracellular bacteria. To prepare the differentiated N2a cells to receive EGC, the cells were washed with 1 ml of prewarmed PBS and suspended in 0.5 ml of CRL-2690 media containing gentamicin at a final concentration of 40 μg/ml. At t= 2.5 h, the infected CRL-2690 cells were detached with Cellstripper (Corning), counted using a hemacytometer, suspended in media with gentamicin, and added to wells containing N2a cells at a 4:1 ratio (EGC:N2a). The 24-well dish was centrifuged at 200 *x g* and then the cells were co-cultured at 37°C with 5% CO_2_.

At the indicated time points, coverslips were washed twice with prewarmed PBS, fixed in 4% buffered paraformaldehyde for 15 min, blocked in 300 µl of antibody buffer (TBS + BSA 1% w/v) for 30 min at RT, and then washed 6X with 400 μl of antibody buffer. Coverslips were stained first with rabbit *Listeria* O antiserum polyclonal (1:100 BD Difco) for 45 min followed by 6X washing and then AF405-conjugated goat anti-rabbit secondary antibody (1:100; Invitrogen) for 45 min at RT for 30-60 minutes. After washing again 6X in antibody buffer, the cells were permeabilized TBS-TX for 30 min at room temperature. Intracellular staining was performed with a cocktail containing 1:40 Texas Red-X phalloidin (ThermoFisher), eFlour-660-conjugated β3-tubulin (1:40; clone 2G10-TB3; Invitrogen) in TBS-TX for 1 h at RT. Finally, the coverslips were washed 5X with TBS-TX, then washed 5X with TBS, dried, and mounted with ProLong Diamond anti-fade (Invitrogen). Images were taken on a Nikon AXR Inverted Confocal Microscope and analyzed in Nikon Element version 6.10.0 and ImageJ version 1.54p.

### Statistical analysis

Statistical analysis was performed with Prism version 10.5.0 (Graph Pad). The specific test used is indicated in each figure legend. Statistical significance is indicated as follows: *, *P*< 0.05; **, *P* < 0.01; ***, *P*< 0.01; ****, *P*<0.001.

## Acknowledgments

The authors thank Tanya Myers-Morales for technical assistance, William Bailey and Dr. John Gensel for consultations in developing the vagus nerve harvest protocol, and Dr. Jamila Tucker for helpful discussions. Figures 2C, 5A and 6A were created in BioRender. Some of this work was supported by the Biospecimen Procurement and Translational Pathology Shared Resource of the University of Kentucky Markey Cancer Center (P30CA177558). This study was funded by a grant from the National Institutes of Health (R01 AI167953) to S.E.F.D.

